# Dose-Dependent Effects of PFOS on Thyroid Hormone-Driven Neurodevelopment and Metamorphic-like change in *Xenopus laevis* larvae

**DOI:** 10.1101/2024.10.30.621080

**Authors:** Katy Zirkle, Ebony D. Myers, Christopher K. Thompson

## Abstract

PFOS is a banned stain repellent that is a type of PFAS, a category of chemicals commonly referred to as “forever chemicals.” These substances are known for their bioaccumulative nature and their resistance to degradation in the environment. Emerging data indicates that PFOS may disrupt thyroid hormone (TH) signaling, which plays a pivotal role in healthy vertebrate development, particularly neural development. Our study investigates whether PFOS interferes with TH-dependent developmental changes, with focus on the developing brain, in *Xenopus laevis* larvae. To assess this, we exposed five-day old larvae to varying concentrations of PFOS (25 μg/L, 2.5 μg/L, and 250 ng/L). Additionally, some larvae were exposed to either 15 μg/L or 1.5 μg/L of thyroxine to evaluate PFOS’s potential interference with TH-dependent developmental mechanisms. After four days, we euthanized larvae and examined body length and hind limb size, dissected out brains and performed immunostaining for markers of neuronal proliferation and apoptosis. We found that PFOS significantly impeded overall body growth in a dose-dependent manner, irrespective of TH status. Moreover, PFOS impaired TH-stimulated hind limb development. In the brain, PFOS had no effect on proliferation but hindered TH-stimulated growth of the optic tectum. Last, PFOS increased apoptosis in the optic tectum, indicative of cytotoxicity. These findings suggest that PFOS may have the capacity to disrupt TH-dependent mechanisms involved in brain development. The observed pattern of change also suggests that its impact is cytotoxic and may not stem solely from a canonical endocrine disruption mechanism.

## Introduction

Polyfluoroalkyl substances (PFAS) are a class of chemicals deemed “forever chemicals” due to their persistence in nature (Allen, 2018). Their exceptionally stable structure coupled with an ability to bioaccumulate poses toxicological risks to life on Earth (Cousins et al., 2020; Wee and Aris, 2023). These compounds are synthetically produced and have served a multitude of industrial applications. PFAS have characteristically high solubility in water, leading to relatively quick and widespread contamination within water sources (Shi et al., 2009; Wee and Aris, 2023; Hohweiler et al., 2024). Before 2003, perfluorooctane sulfonate (PFOS) was the main ingredient in Scotchgard, a fabric protector made by 3M; it is now one of the most prevalent strains of PFAS detected in the environment. PFOS exposure is ubiquitous; one study found that more than 98% of 2,094 blood samples were positive for PFOS (Calafat et al., 2007). The combined effects of exposure to different PFAS types, such as PFOS and PFHxS, are complex; research indicates that the toxicity of mixtures does not necessarily follow simple additive models, as demonstrated in amphibian models, where combined PFAS exposure altered growth patterns and delayed development beyond what individual exposures predicted (Hoskins et al., 2022). Research suggests that exposure to PFAS, including PFOS, can have detrimental health effects in both plants and animals. PFAS chemicals may also have effects on humans; including potential developmental toxicity in children, carcinogenicity, disruption of metabolic function in adults, and endocrine disruption-like effects on hormones such as thyroid hormone (TH) (Kumari and Shahbaz, 2024; Nannaware et al., 2024). Consequently, PFOS was added to the list of chemicals to be restricted by the Stockholm Convention in 2009 (Paul et al., 2009; Ahrens and Bundschuh, 2014). Regulation of the production of PFOS is complicated by the prevalence of already existing PFOS materials that will continue to emit pollutants throughout their lifecycle as well as a lack of a global standard for production and use of these materials (Cousins et al., 2020).

Currently, there is a loose constellation of observations suggesting that exposure to PFAS disrupts the regulation and function of TH-signaling, although the specific mechanism by which this occurs is not entirely clear and likely varies depending upon the type of PFAS compound (Coperchini et al., 2017a, 2021a). The thyroid gland primarily synthesizes thyroxine (T_4_), which circulates around the body and is locally converted to triiodothyronine (T_3_), whereupon it acts on the thyroid hormone receptors TRα and TRβ to induce changes in gene expression. While TH is well known for its role in metabolic processes in adults, it regulates key aspects of vertebrate development, particularly the developing brain (Zoeller and Rovet, 2004; Rovet, 2014). Chemicals that disrupt the functioning of these hormones have high potential to disrupt development, with the possibility of life-long detrimental effects. Studies performed on mice and rats show that exposure to high doses of PFOS during pregnancy lead to significant decreases in neonate survival rates and slowed development in the pups that persisted to adulthood (Lau et al., 2003). The higher mortality rates and delayed growth were also coupled with a decrease in free T4 in the rats, which may suggest PFOS’s mechanism targets hypothalamus-pituitary-thyroid (HPT) axis (Lau et al., 2003). Another study found that PFOS exposure in zebrafish induced a number of changes in TH-signaling and concluded that PFOS disrupts thyroid hormone regulation via both direct effects on the thyroid gland and indirect effects on the regulation of the HPT axis (Shi et al., 2009). Notably, there are two observations indicating that PFOS exposure increases expression of TRβ in fish, which suggests that PFOS exposure results in enhanced TH-signaling (Shi et al., 2009; Rodríguez-Jorquera et al., 2019). Last, there is some limited evidence, including modeling, that suggests that some PFAS have the capacity to bind to TR and may fit into the binding pocket (Ren et al., 2015; Mortensen et al., 2020; Young et al., 2021). Overall, these findings underscore the potential of PFOS to significantly interfere with thyroid hormone signaling, and therefore development, across different species.

*Xenopus laevis*, the African clawed frog, provides unique advantages for observing dysfunction in T_4_-driven processes. Amphibian metamorphosis is particularly sensitive to changes in thyroid hormones because it is utterly dependent upon a surge in circulating TH (Tata, 1999; Brown and Cai, 2007; Buchholz, 2015, 2017). A study designed to model how PFOS disrupts normal thyroid function in *X. laevis* found significant increases in TrβA, BTEB, and DI2 mRNA expression in larvae exposed to environmentally relevant concentrations of the chemical. All three of these genes are TH sensitive and play roles in regular thyroid development and activity, with some of their results suggesting PFOS acts to mimic and therefore potentially disrupt normal thyroid hormone physiology (Cheng et al., 2011). The action of thyroid hormone agonists and antagonists can be monitored in tadpole development within a relatively short period due to changes in TH-dependent gene expression (Zhang et al., 2006). Larvae exposed to T4 will typically demonstrate enhanced cell proliferation in the brain and precocious hindlimb development (Brown et al., 2005; Thompson and Cline, 2016). Exposure to a T4 agonist (or a RXR agonist, see discussion) typically results in similar effects (Veldhoen et al., 2006; Zhang et al., 2006; Mengeling et al., 2016). Furthermore, the hypothalamic-pituitary-thyroid (HPT) axis in amphibians is particularly sensitive to changing stimuli. For instance, environmental stressors such as limited food availability or increased water pollution activate the hypothalamic–pituitary–adrenal axis to release corticotropin-releasing hormone which, in turn, activates the production and release of thyroid-stimulating-hormone, ultimately speeding up the rate of metamorphosis in *Xenopus* (Rousseau et al., 2021). TR antagonists, on the other hand, should impose opposite outcomes in developing larvae and tadpoles and therefore would result in limited morphological changes that are typically associated with TH and there would be no increases expression of TH responsive genes (Lim et al., 2002; Opitz et al., 2006b, 2006a).

Our study investigates the relationship between PFOS exposure and TH-dependent development in *Xenopus laevis* larvae. We euthanized *Xenopus* that had been exposed to varying concentrations of PFOS and T_4_ or a combination of the two at five days post-fertilization. Following euthanasia four days later, the body lengths and hind limb sizes of larvae were measured to identify the potential effects of PFOS on TH-mediated changes in development. We also examined the effects of PFOS in dissected and immunostained brains, with staining for a marker of neuronal proliferation in some brains and in others for two different markers of apoptosis. Our findings suggest PFOS has the capacity to interfere with TH-dependent mechanisms of development, including in the developing brain, but that PFOS does not appear to act as a specific TR agonist or antagonist and is instead cytotoxic. These results provide the groundwork for further investigation into the mechanisms by which PFOS affects neurological functioning and brain development.

## Materials and methods

### Animals

We used *Xenopus laevis* albino larvae for this experiment, having been raised in the lab after an HCG-stimulated breeding with mature male and female albino frogs. Fertilized eggs were held in a 20-gallon tank for five days at 20° C. Then, stage 44 larvae were transferred into treatment bowls made of HDPE plastic (to prevent sorption of PFOS out of solution) and moved into a 22° C incubator for four days.

### PFOS and thyroxine treatment

Granulated PFOS (TRC Inc; H268700) was diluted into deionized water (250 mg/L), diluted again 1:10 in water and adjusted for pH (7.0), then diluted 1:10 in Steinburg’s solution up to three more times to create stock solution for each PFOS treatment. 2 ml of stock PFOS solution was added to 198 ml of either Steinburg’s or thyroxine to a final concentration of 25 ug/L, 2.5 ug/L, or 250 ng/L. Granulated thyroxine (Sigma; T2376) was diluted into 50 mM NaOH, then serially diluted 1:10 into Steinberg’s solution until final concentration (15 ug/L or 1.5 ug/L). Once final solutions were made, we placed *Xenopus* (n = 72-93 in each group) into 200 ml of one of 12 treatments and assessed as three separate subgroups with one control and three different doses of PFOS (Fig 1).

**Figure 1.**
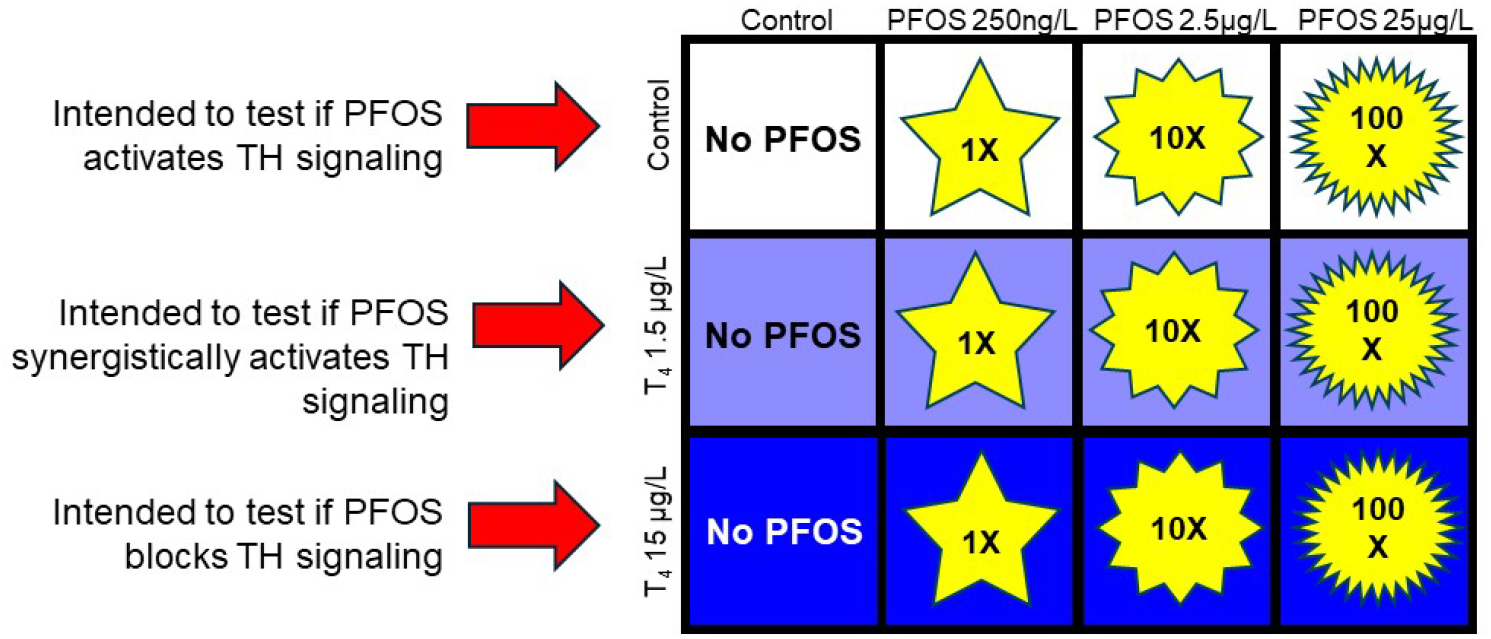
Experimental setup for assessing PFOS effects on thyroid hormone (TH) signaling in Xenopus laevis larvae. Larvae were divided into 12 groups across three conditions: without T_4_ (top row), with low T_4_ (1.5 μg/L, middle row), and with high T_4_ (15 μg/L, bottom row). Each condition was further divided into four PFOS treatment levels: no PFOS (control), 250 ng/L, 2.5 μg/L, and 25 μg/L. The figure shows increasing PFOS concentrations from left to right, with yellow stars indicating PFOS presence. This arrangement aims to determine whether PFOS activates, synergistically enhances, or blocks TH signaling.

### Sacrifice and tissue fixation

After four days of treatment, larvae were killed with an overdose of 0.4% MS-222 and fixed in 4% paraformaldehyde for at least 24 h.

### Body length

42-54 larvae were selected at random for body length analysis. Entire bodies of treated larvae were imaged using a Nikon stereomicroscope (1X) (Fig 2A). Body length was measured from snout to tail tip using ImageJ. The observer was blind to treatment.

**Figure 2.**
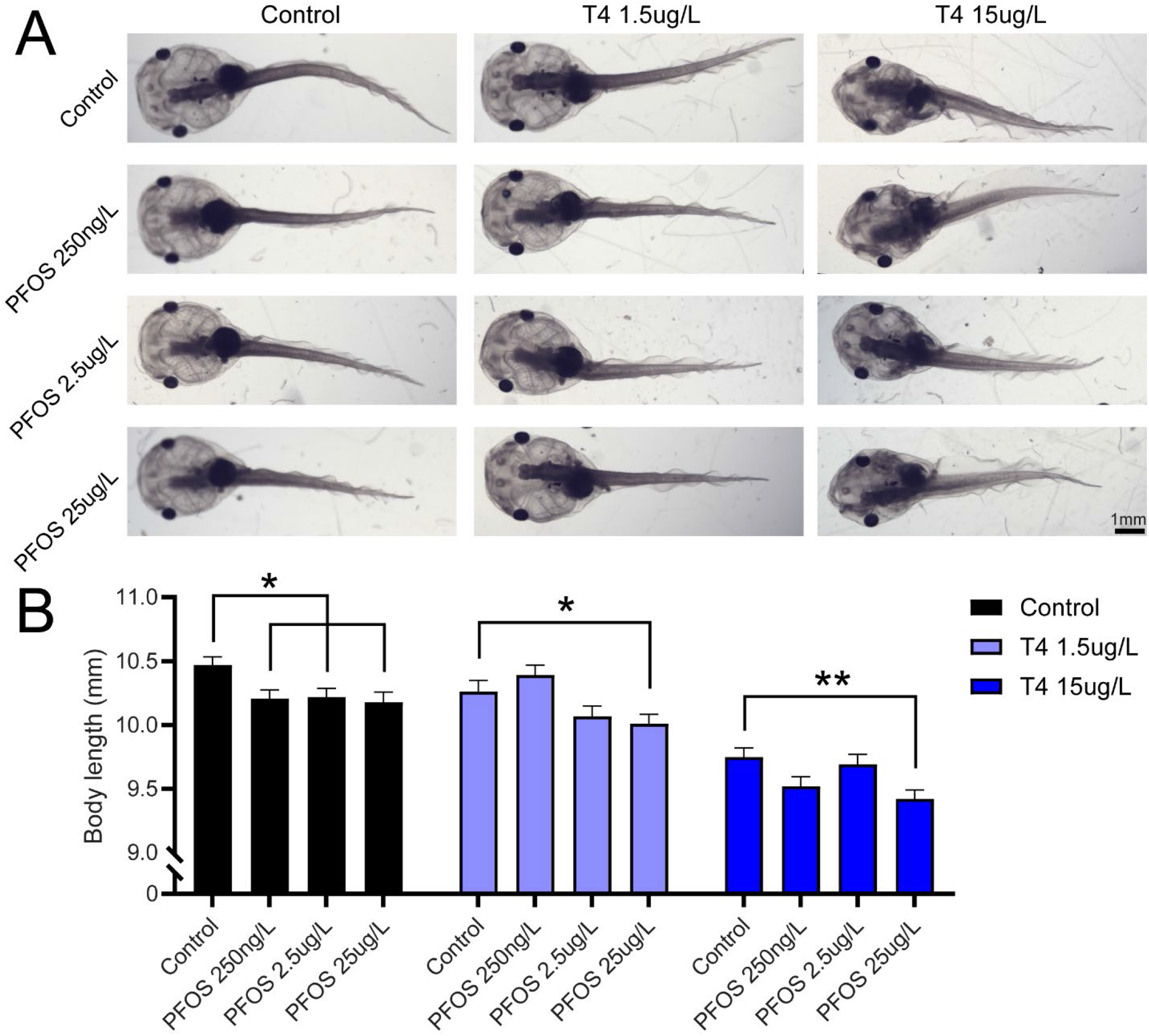
PFOS impaired growth of Xenopus body length. A) Representative images of tadpoles in treatment groups. B). Quantification of body length in Xenopus Larvae treated with either CNTL bath or thyroxine (T4,1.5 µg/L or 15 µg/L) and/or treated with PFOS (250ng/L, 2.5 µg/L, or 25 µg/L) for 4 days. PFOS treatment impaired growth of body length in CNTL and T4-treated larvae. * = p < 0.05, ** = p < 0.01

### Hind limb

10-12 larvae were selected at random for hind limb analysis. We imaged the right side of whole larvae using a Nikon stereomicroscope (5X) and measured the area of the hind limb using the perimeter of the oblique side of the limb bud with ImageJ (Fig 3A and B). The observer was blind to treatment.

**Figure 3.**
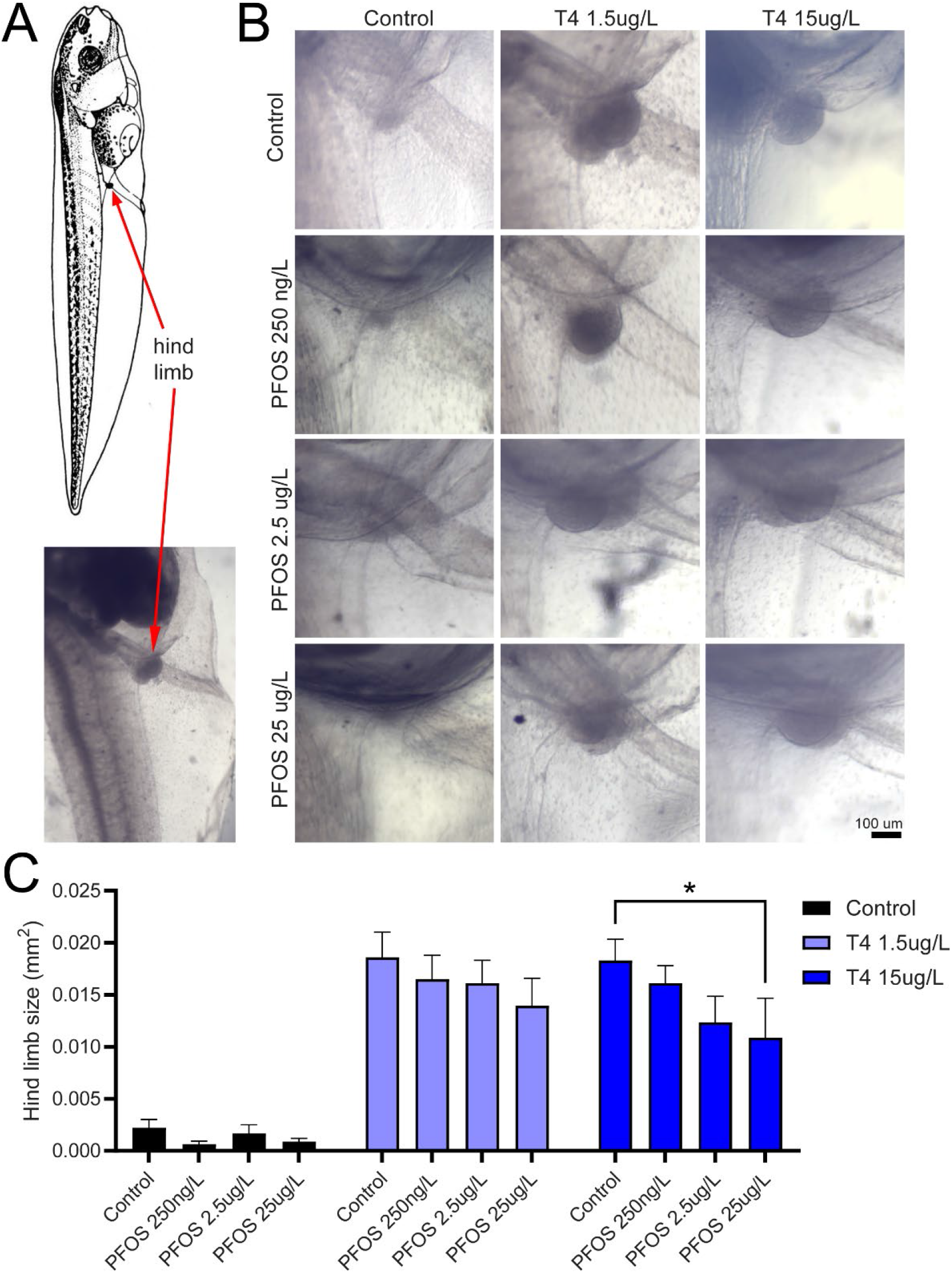
PFOS impaired TH-mediated growth of the hindlimb. A) Right side of a Xenopus laevis larvae, illustrating the hind limb. Bottom is an image of the location of the hind limb in a larva treated with T4. B) Representative images of hind limbs in treatment groups. C) Quantification of hind limb size in Xenopus larvae treated with either CNTL bath or thyroxine (T4,1.5 µg/L or 15 µg/L) and/or treated with PFOS (250ng/L, 2.5 µg/L, or 25 µg/L) for 4 days. PFOS 25 µg/L impaired hind limb growth in larvae treated with T4 15 µg/L. * = p < 0.05

### Brain dissection

After fixation, we dissected brains from a subset of larvae using micro-forceps and placed them into microcentrifuge tubes containing PBS-TX (triton X 100, 0.1%) until processed for immunostaining.

### Immunostaining

To prepare brains for measurement of proliferation and morphological changes in the optic tectum (Fig 4A), we washed brains (n = 23-25 from each group) in PBS-TX, placed them in blocking buffer (2.5% normal goat serum in PBS-TX) for 1 h, and then incubated them overnight in a primary antibody for phospho-Histone 3 (pH3, Millopore-Sigma, H0412, made in rabbit, 1:1000) in blocking buffer at 4°C with gentle rotation. Brains were then washed in PBS-TX and incubated with an Alexa Fluor 488 (1:400, Life Technologies, A11094, goat anti-rabbit) with gentle rotation at room temperature for 4 h. We then counterstained brains for 15 min with Sytox-O (1:500 in PBS, ThermoFisher, S34861), a nuclear marker. We then washed them in PBS and coverslipped them using 6M urea in 50% glycerol as a mounting medium.

**Figure 4.**
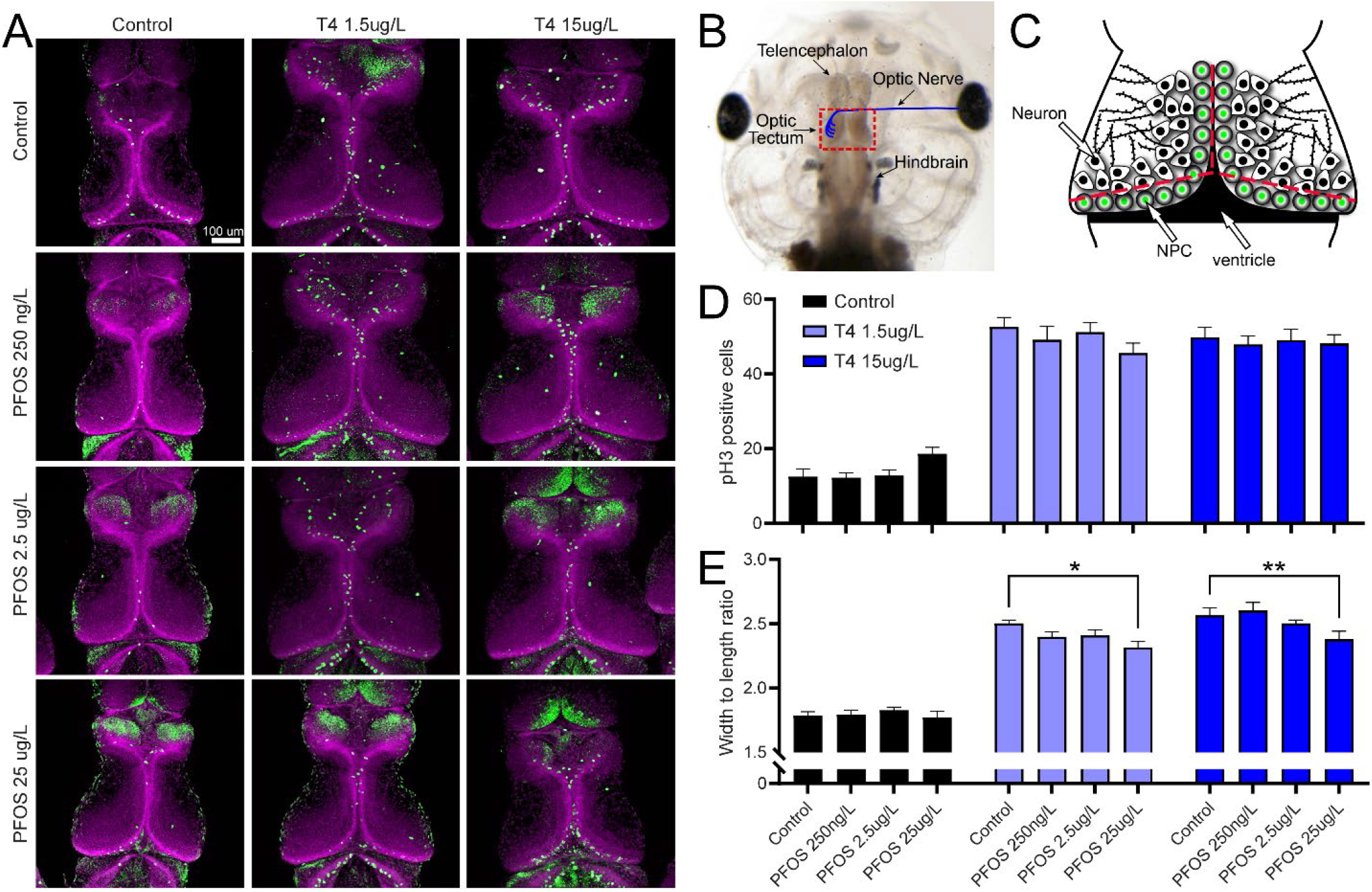
PFOS impaired TH-mediated changes in Xenopus optic tectum morphology. A) Example brains of *Xenopus laevis* larvae immunostained to allow for quantification of proliferation and brain morphology. green: phosphorylated histone 3 (pH3); magenta: Sytox-O, a nuclear marker used as a counterstain to reveal brain morphology. Xenopus were treated with either CNTL bath or thyroxine (T4,1.5 µg/L or 15 µg/L) and/or treated with PFOS (250ng/L, 2.5 µg/L, or 25 µg/L) for 4 days. B) Image of a *Xenopus* head illustrating the major subdivisions of the brain. C) Diagram of the optic tectum (inset in B) illustrating the ventricular zone lined with neural progenitor cells (NPC) that divide and differentiate into neurons. The dotted lines illustrate the anatomical features used to calculate width and length of the optic tectum. D) PFOS had, at best, a marginal effect on proliferation, with a slight dose-dependent increase observed in CNTL brains and a slight dose-dependent decrease observed in T4-treated brains. No statistically significant effects were observed. E) PFOS treatment impaired the T4-induced increase in width to length ratio is a dose-dependent manner. * = p < 0.05, ** = p < 0.01

To stain brains for apoptotic markers (Fig 5A), we stained brains (n =12-13 per group) using the same procedure as above, except that brains were incubated overnight in apoptosis-related primary antibodies (H2A.X, 1:500, EMD Millipore, 05-636, made in mouse; and caspase 3, 1:500, abCam, AB13847, made in rabbit) overnight in blocking buffer at 4°C with gentle rotation. The next day, brains were incubated in the appropriate secondary antibody (h2ax with Alexa Fluor 488, 1:400, Life Technologies, A11029, goat anti-mouse; caspase-3 with Alexa Fluor 647, 1:400, Life Technologies, A21244, goat anti-rabbit) for 4 h, washed them again in PBS-TX, and incubated them in Sytox-O. Brains were washed in PBS and cover-slipped in a well on a slide with mounting medium.

**Figure 5.**
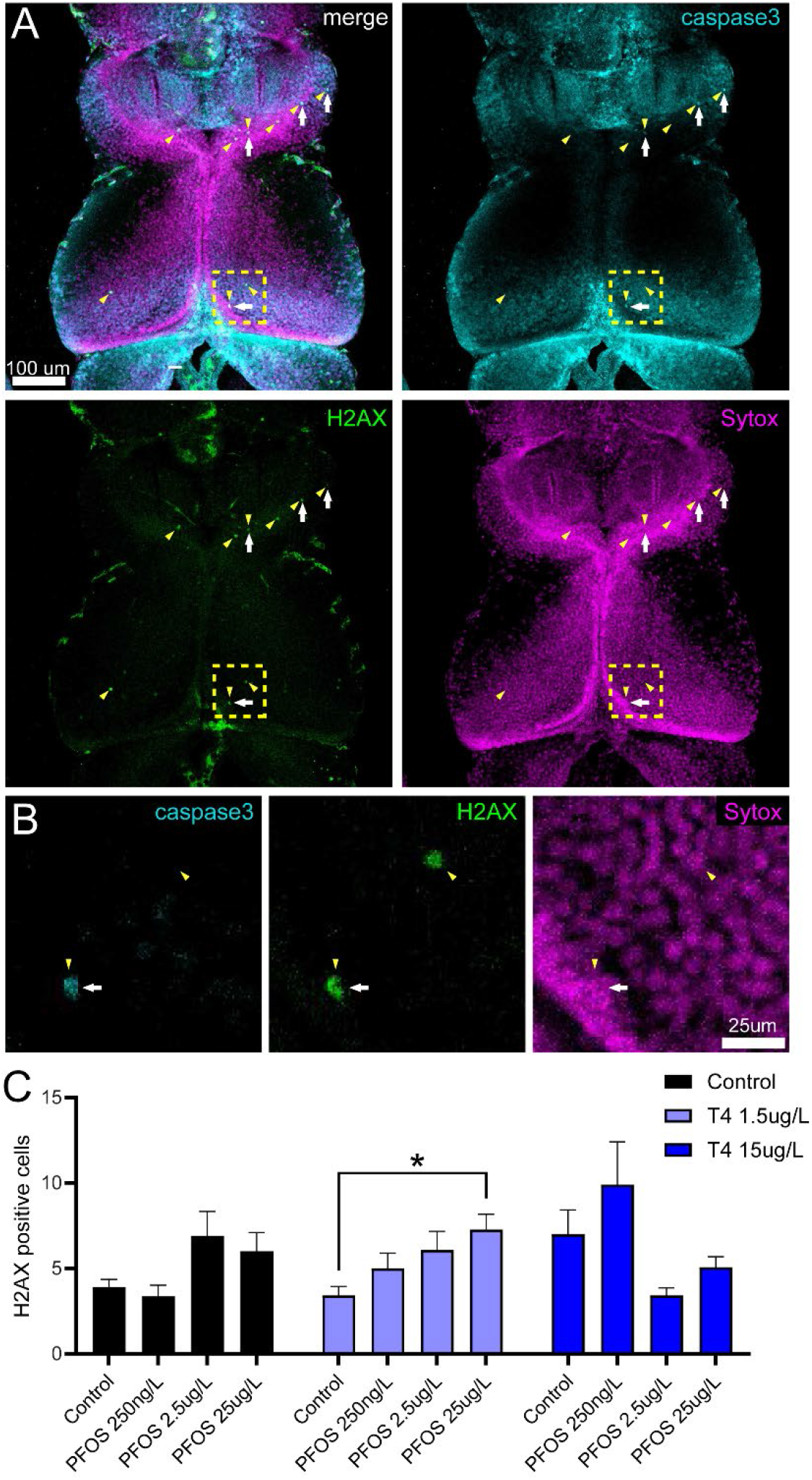
PFOS induced an increase in cell death in the Xenopus optic tectum. A) Brains of *Xenopus laevis* larvae immunostained to allow for quantification of programmed cell death. Green (arrowheads): Anti-phospho-Histone H2A.X (H2AX); cyan (arrows): activated caspase-3; magenta: Sytox-O, a nuclear marker. Brains were treated with either CNTL bath or thyroxine (T4,1.5 µg/L or 15 µg/L) and/or treated with PFOS (250ng/L, 2.5 µg/L, or 25 µg/L) for 4 days. B) Zoomed in view of the inset in A illustrating caspase 3 and H2A.X positive cells. C) PFOS treatment resulted in a significant increase in the number of H2A.X positive cells in tecta of *Xenopus* treated with T4 (1.5 µg/L) in an apparent dose-dependent manner. * = p < 0.05

### Imaging and image analysis

Immunostained brains were imaged on a Leica SP8 confocal microscope. The same imaging parameters were used for all brains in all groups in each set of immunostained brains. We imaged starting from the dorsal side of the optic tectum moving ventrally for 70 optical sections (72.8 µm) in all brains.

All quantification was done in ImageJ. We counted all pH3+ cells that were in the proliferative zone adjacent to the ventricle; most, if not all, or those cells are presumed to be neural progenitor cells which divide and give rise to neurons (Fig 4B) (Bestman et al., 2012). Changes in optic tectum morphology were assessed by measuring the width and the length of the tectum from a z-projection of the confocal stack (Fig 4C). We used the width/length ratio as a measure of change in tectum morphology because it is a more sensitive measure of the effects of T3 than total volume (Thompson and Cline, 2016). For apoptosis, we observed more H2A.X+ cells than caspase-3+ cells. Nearly every caspase-3+ cell was also H2A.X+ (Fig 5B), so we elected to quantify just the H2A.X+ cells in the entire optic tectum.

### Statistical analysis

Two-way ANOVA’s and subsequent graphs were constructed using Prism. Dunnett’s multiple comparisons test were used post-hoc to determine significant differences of PFOS treated groups relative to the subgroup control group.

## Results

Our experimental outcomes reveal PFOS exposure had a significant impact on several aspects of the development of *Xenopus laevis* larvae, including impairment of several aspects of thyroid hormone-mediated changes in development, particularly neural development.

### Four days of PFOS treatment had no effect on mortality

We monitored the number of dead *Xenopus* in each treatment group each day. We found that there was no significant increase in mortality in any of our treatment groups, with no more than 1-2 dead larvae observed in each group per day. Most days, there were no dead larvae observed in each group.

### PFOS treatment resulted in shorter larvae

If PFOS stimulates TH-signaling, resulting in metamorphic-like changes in development, then we would expect PFOS treatment in pre-metamorphic larvae to result in shorter body length due to the initiation of tail resorption mechanisms. We found that exposure to PFOS significantly impaired the growth of body length in control and thyroxine-treated *Xenopus*. The most pronounced reduction in growth was observed in larvae treated with the highest concentration of PFOS (25 μg/L), which showed significant impairments (*p < 0.05, **p < 0.01) across both control and thyroxine-treated groups. Furthermore, all doses of PFOS resulted in shorter body lengths relative to control bath treatment. This result is consistent with the hypothesis that PFOS acts to stimulate TH-like tail resorption. The result is also consistent with the hypothesis that PFOS inhibits overall developmental growth and is generally cytotoxic.

### PFOS treatment impaired TH-stimulated growth of the hindlimb

If the effects of PFOS on body length were due to stimulation of TH-signaling, then PFOS treatment should likewise stimulate premature growth of the hindlimb. If the effects of PFOS on body length were due to cytotoxic mechanisms that impair overall growth of tissues, then PFOS treatment should instead impair growth of the hindlimb. We found that the development of hind limbs was notably stunted in the presence of PFOS, especially at the highest concentration (25 μg/L) and in combination with the highest dose of T4 (15 μg/L) (Figure 2C). This result is consistent with the hypothesis that PFOS induces dose-dependent toxicological effects on morphological development, including the ability to impair TH-stimulated growth of the hindlimb. It also is inconsistent with the hypothesis that PFOS itself has the capacity to stimulate TH-like mechanisms to stimulate growth of the hindlimb.

### PFOS treatment had no effect on proliferation but impaired TH-stimulated changes in morphology of the optic tectum

We also examined the effects of PFOS on brain development by analyzing its impact on the optic tectum. We found that T4 treatment resulted in an increase in cellular proliferation (number of pH3+ cells) in the ventricular zone and in an increase in the width-to-length ratio of the optic tectum, a phenomenon known to occur in pre-metamorphic *Xenopus* (Thompson and Cline, 2016). In contrast, we found that PFOS treatment had no significant impact on proliferation. In control conditions, a slight, but non-significant dose-dependent increase in neuronal proliferation was observed with increasing concentrations of PFOS. Conversely, in thyroxine-treated brains, PFOS exposure led to a slight decrease in proliferation, which was also dose-dependent (Figure 3B). Additionally, PFOS exposure altered the T4-induced changes in brain morphology, specifically impacting the width-to-length ratio (Figure 3C). These results are consistent with the conclusion that PFOS impairs developmental growth mechanisms but does not appear to stimulate TH-signaling.

### PFOS increased apoptosis in TH-treated larvae brains

Given the above results suggest that PFOS impairs developmental growth mechanisms, this suggests that PFOS may be cytotoxic and may induce programmed cell death, or apoptosis. To test this, we immunostained 10-12 prematamorphic *Xenopus* brains for two markers of cell death, activated caspase 3 and phosphorylated H2A.X (Figure 4A). We found that apoptosis was relatively low in all *Xenopus* tecta but that there were more visible H2A.X+ cells than caspase 3+ cells, although virtually every single caspase-3+ was likewise positive for H2A.X (Figure 4B). Thus, we elected to quantify just H2A.X+ cells. We found that PFOS had a dose-dependent effect on cell death in *Xenopus* treated with 1.5 μg/L T4, with the highest concentration of PFOS resulting in a significant increase in apoptosis relative to control (Figure 4C). This result underscores the potential cytotoxic effects of PFOS, especially in the face of growth mechanisms stimulated by TH.

## Discussion

PFOS has emerged as a significant environmental contaminant with potential endocrine-disrupting effects, particularly on TH signaling. TH plays a critical role in regulating growth, metabolism, and developmental processes in all vertebrates, especially in brain development (Zoeller and Rovet, 2004; Rovet, 2014). In amphibians like *Xenopus laevis*, TH plays a major role in development as the key driver of metamorphosis. If PFOS has the capacity to interfere with TH-mediated metamorphosis by mimicking or inhibiting the action of TH, then *Xenopus* would be an ideal animal model to evaluate its capacity for endocrine disruption. One major advantage amphibian metamorphosis has is that it includes a constellation of developmental changes that reflect both growth and regression. For example, a compound that initiates TH-signaling will result in the regression of the tail via apoptosis but also growth the hind limbs. In the brain, it will result in growth of the optic tectum and increase in neuronal proliferation but also regression of the telencephalon (Thompson and Cline, 2016). In contrast, a compound that antagonizes TH-signaling should do the exact opposite when TH is present (e.g. continual increase in body length but also diminished or no growth of the hind limb). It is an ideal animal model to disentangle these effects from a compound that is simply cytotoxic and that does not interfere directly with TH-signaling mechanisms because a cytotoxic compound will typically result in hindering developmental growth of all kinds. PFOS appears to be largely cytotoxic, at least with respect to the aspects of the TH system evaluated in this study.

Our results reveal that PFOS exposure significantly impacts the development of *Xenopus laevis* larvae, including disruption of TH-mediated changes in neural development. Although PFOS treatment for four days did not affect mortality rates, it did result in shorter larvae, which would be consistent with the hypothesis that PFOS initiates tail resorption via a TH-mediated mechanism. In contrast, PFOS impaired TH-stimulated hindlimb growth, which runs contrary to that hypothesis. In the optic tectum, PFOS treatment did not affect proliferation but impaired TH-stimulated morphological increase in optic tectum size, similar to the hind limb observations. Additionally, PFOS increased apoptosis in TH-treated larval brains, underscoring its cytotoxic potential. Overall, these findings indicate that PFOS disrupts developmental processes in *Xenopus laevis* larvae, affecting growth mechanisms and causing neurotoxicity without acting as a canonical TH disruptor that acts upon TR.

Our study is primarily designed to test if PFOS exposure (directly or indirectly) interferes with how TH acts on TRs. There are two types of TRs: TRα, which is the first receptor to express (even before thyroid gland development) and TRβ, whose expression is directly correlated with TH concentrations (Yaoita and Brown, 1990; Sachs et al., 2000; Schreiber et al., 2001). Ligand-binding to TRs initiate substantial change in gene expression in TH-sensitive tissues because TRs act as transcription factors (Brent, 2012). Furthermore, TRs bind to TH response elements as a heterodimer, with retinoic acid-X receptor (RXR) acting as a binding partner (Mangelsdorf and Evans, 1995). Thus, TH signaling can be modified by RXR agonists and antagonists; indeed, there is a growing body of evidence that agonists of RXR result in enhancement of TH signaling developing *Xenopus* (Mengeling and Furlow, 2015; Mengeling et al., 2016, 2018, 2022). Our second subgroup (consisting of *Xenopus* treated with T4 at 1.5 ug/L) is intended to determine if PFOS has the capacity to work synergistically with a relatively low concentration of T4 to initiate metamorphic-like change, much like a putative RXR agonist should act. The only evidence we observed that would be consistent with the hypothesis that PFOS acts as an RXR agonist is that increasing concentrations of PFOS with T4 (1.5 ug/L) resulted in shorter larvae, but that result is also consistent with the hypothesis that PFOS is cytotoxic and impairs developmental growth of all kinds. In contrast, PFOS with T4 (1.5 ug/L) resulted in a marginal decline in TH-mediated increase in optic tectum size and hind limb size, which runs contrary to the hypothesis that PFOS may as a RXR agonist but is consistent with the cytotoxicity hypothesis. Thus, we conclude that PFOS does not act synergistically with TH by acting on RXR to enhance TH-signaling.

We observed that PFOS exposure results in impaired growth of various tissues. These observations are consistent with experimental evidence from other studies, including two studies that likewise observed reduced body length in PFOS-treated fish (Shi et al., 2009; Rodríguez-Jorquera et al., 2019). Similar impacts on fetal growth have been observed in mice and rats (Grasty et al., 2003; Lau et al., 2003, 2006; Thibodeaux et al., 2003; Luebker et al., 2005; Yahia et al., 2008). A correlation between maternal exposure and low birth weight have been observed in humans in serval studies as well (Apelberg et al., 2007; Monroy et al., 2008; Fei et al., 2009). It is important to note that substantially higher serum concentrations of PFOS and PFOA required to observe toxic effects in animals compared to humans, which weakens the biological plausibility of a direct causal relationship in humans based on a recent toxicological meta-analysis (Negri et al., 2017). It may be that the exposures in humans that lead to decrease in birthweight are dependent on exposure to other PFAS or other chemicals as a mixture (Gundacker et al., 2022). Evidence suggests that a mixture of PFAS compounds may act in unexpected ways, deviating from predicted additive outcomes and altering expectations about dose-dependent consequences (Hoskins et al., 2022). These findings underscore the complexity of PFAS exposure effects and the need for further research into the combined impacts of multiple environmental contaminants on human health.

Our results suggest that PFOS exposure does not directly disrupt some aspects of TH-signaling as a canonical endocrine disruptor, particularly with respect to agonism and antagonism of TRs, despite the fact that other studies suggests that some PFAS, including PFOS may bind to TR as an agonist (Ren et al., 2015; Mortensen et al., 2020; Young et al., 2021). There is some evidence that PFOS has the capacity to disrupt TH in ways that are not addressed by our experiments (Coperchini et al., 2017b, 2021b, 2024). For instance, thyroid peroxidase (TPO) is an enzyme crucial for the synthesis of thyroid hormones, catalyzing the iodination of tyrosine residues in thyroglobulin and the coupling of iodotyrosines to form T3 and T4. Several studies have found that PFOS inhibits TPO activity are other mechanisms by which PFOS PFOS and other perfluoroalkyl substances (PFASs) were found to compete with thyroxine (T4) for binding to transthyretin (TTR), a major thyroid hormone transport protein in the blood. This competitive binding can lead to a reduction in T4 availability for physiological functions (Weiss et al., 2009; Hamers et al., 2020; Langberg et al., 2024). The disruption caused by PFOS could stem from its ability to interfere with the synthesis, release, transport, and receptor binding of TH, ultimately leading to imbalanced hormonal regulation and impaired developmental outcomes.

Overall, these results show that PFOS exposure disrupts developmental processes in *Xenopus laevis* larvae, particular mechanisms of growth. Our results indicate that PFOS does not appear to act as a canonical endocrine disrupting agonist/antagonist that specifically affects thyroid hormone receptor activation in a specific direction. PFOS does appear to have the capacity to interfere with TH-mediated changes in development, however, particularly when TH stimulates growth mechanisms. Last, our data show that PFOS can cause neurotoxicity in the developing brain but did not appear to affect proliferation in the midbrain.

